# MMRi62 induces iron depletion–driven apoptosis through a ferritin-independent mechanism

**DOI:** 10.64898/2026.09.22.753515

**Authors:** F Segui, M Pagnuzzi, J Antiq, J Durivault, V Vial, N Bogliotti, I Leray, M Vucetic, V Picco

## Abstract

The modulation of iron metabolism is increasingly explored as a therapeutic strategy in cancer, particularly through the induction of ferroptosis. Given the central role of ferritin as a major intracellular iron buffer, targeting ferritin represents an attractive strategy to disrupt iron homeostasis; however, the lack of pharmacological approaches capable of selectively targeting ferritin currently limits the therapeutic exploitation of this vulnerability. MMRi62, initially developed to target the MDM2–MDM4 axis, has been proposed to induce ferroptosis *via* ferritin degradation. Here, we revisited this hypothesis in the context of medulloblastoma (MB). Contrary to this proposed mechanism, we found that MMRi62 does not trigger ferroptosis but instead induces robust apoptotic cell death. This effect is observed in both p53-mutant DAOY and p53 wild-type HD-MB03 cells, as well as in c-Myc/OTX2-driven medulloblastoma-like tumours genetically induced in brain organoids. Although ferritin degradation occurs upon treatment, genetic dissection demonstrates that neither ferritin itself nor ferritinophagy are required for MMRi62-induced cytotoxicity. Together, these findings rule out ferritin-dependent mechanisms as primary drivers of cytotoxicity. Instead, we uncovered that MMi62 induces a profound rewiring of iron metabolism consistent with a canonical iron starvation response, accompanied by a marked reduction in intracellular iron levels. Consistently, the UV–visible spectroscopic data were in agreement with the proposed iron-chelating properties of MMRi62. Taken together, our findings identify iron depletion–driven apoptosis, rather than ferroptosis, independently of ferritin degradation, as the primary mechanism of MMRi62 cytotoxicity in MB, refining its mode of action and highlighting iron homeostasis as a therapeutic vulnerability.

## Introduction

Medulloblastoma (MB) is the most common malignant brain tumor in children and continues to be associated with substantial morbidity, largely due to the long-term toxicity of current treatments and the frequent emergence of therapeutic resistance (1,2). These limitations underscore the urgent need to identify exploitable biological vulnerabilities that could be leveraged for more effective and less deleterious therapeutic strategies. Among these, iron metabolism has recently emerged as a critical determinant of tumor cell fitness (3–5). Iron is indispensable for fundamental cellular processes, including DNA synthesis, mitochondrial respiration, and redox regulation. However, its intracellular levels must be tightly controlled, as excess labile iron can catalyse the formation of reactive oxygen species (ROS) and promote oxidative damage, ultimately triggering ferroptosis (6). To cope with this duality, cancer cells actively rewire iron homeostasis pathways, notably through upregulation of ferritin, a key iron storage complex that buffers intracellular iron and limits redox stress (7,8).

Building on this concept, we previously demonstrated that genetic ablation of ferritin heavy chain (FTH) in MB models profoundly sensitizes tumor cells to iron-dependent stresses (9). These findings position ferritin not merely as a passive iron storage protein, but as a central regulator of tumor adaptation to redox and iron imbalance, and therefore as a potential therapeutic vulnerability. Accordingly, targeting ferritinophagy has emerged as a promising therapeutic strategy, particularly in cancers characterized by altered iron homeostasis and increased iron dependency (reviewed in (10)). In particular, enhancing nuclear receptor coactivator 4 (NCOA4)-mediated ferritinophagy has been extensively investigated given the central role of NCOA4 in delivering ferritin to lysosomes for degradation, thereby promoting iron release and potentially sensitizing cancer cells to iron-dependent cell death (11,12). To date, strategies targeting ferritin have relied predominantly on genetic approaches, which are not readily translatable to the clinic. Pharmacological strategies remain poorly developed, with limited evidence regarding their clinical feasibility, particularly in brain cancers. In this context, MMRi62 has recently attracted attention. Initially developed as an inhibitor of the MDM2– MDM4/p53 axis (13), this compound has also been reported to induce ferritin degradation and trigger ferroptosis in pancreatic cancer models, raising the possibility of exploiting ferritin-dependent cancer vulnerabilities pharmacologically (14). Yet, this proposed mechanism appears at odds with a broader body of work on MMRi compounds, including MMRi62 itself, which predominantly induce apoptotic cell death through modulation of the MDM2–MDM4 E3 ubiquitin ligase complex, often independently of p53 status (15).

Here, using two MB cell lines with distinct p53 status we show that MMRi62 induces apoptosis rather than ferroptosis, independently of p53 status. Although MMRi62 treatment was accompanied by ferritin degradation, genetic deletion of *FTH1* or *NCOA4* did not prevent or substantially alter MMRi62-induced cell death, indicating that disruption of the ferritin– NCOA4 axis is not required for its cytotoxic activity. Instead, our findings indicate that the cellular effects of MMRi62, including ferritin degradation, largely stem from its iron starvation induced properties. Together, these results identify iron chelation-driven apoptosis as the predominant mechanism of MMRi62 cytotoxicity in MB cells and challenge the notion that ferritin degradation and ferroptosis are its primary mode of action. Nevertheless, the iron-chelating properties of MMRi62 and their potential therapeutic relevance in medulloblastoma warrant further investigation.

## Materials and Methods

### Cell lines and culture conditions

Human Medulloblastoma (MB) DAOY and HD-MB03 cells were obtained from the American Type Cancer Collection (ATCC, Manassas, VA, USA) and authenticated in 2026 (Eurofins Genomics, France). They were routinely tested for Mycoplasma (PlasmoTest Mycoplasma Detection Kit; InvivoGen) and cultivated up to a 10^th^ passage. Cells were grown at 37 °C/5% CO_2_ in DMEM (Gibco, Thermo Fisher Scientific Inc, MA, USA) supplemented with 8% Gibco™ Fetal Bovine Serum (FBS, certified, heat inactivated, Thermo Fisher Scientific Inc, MA, USA).

### Genetic deletion of ferritin heavy chain (FTH) and nuclear receptor coactivator 4 (NCOA4) using CRISPR-Cas9

DAOY and HD-MB03 wildtype (WT) cells were transfected using jetPEI (Polyplus, 101000053) according to the manufacturers instructions with pSpCas9(BB)-2A-GFP (PX458) plasmid (a gift from Feng Zhang; Addgene plasmid #48138; http://n2t.net/addgene:48138; research re-source identifier: Addgene_48138) (16) containing clustered regularly interspaced short palindromic repeats (CRISPR)-CRISPR-associated protein 9 (Cas9) targeting the following regions: *FTH1* exon 1 immediate flanking region gRNA (5′- **G**TTACCTGTCCATGgtgagcg - 3′), and *FTH1* exon 3 gRNA (5′- gggttccaatactcacATGG - 3′) or *NCOA4* exon 2 : 5-GTGAGGTGTAGTGATGCACGG-3 and NCOA4 exon 3 : 5- GTCTTAGAAGCCGTGAGGTA-3. Bold **G** in 5 is added due to the transcription initiation requirement of a G base for human U6 promoter. GFP-positive cells were single-cell sorted by flow cytometry 24 h post-transfection into 96-well plates containing DMEM supplemented with 8% FBS. Knockout clones were screened by immunoblotting and validated by Sanger sequencing (Eurofins Genomics, France). Two independent knockout clones of each cell lines were selected for experiments, to minimize clonal effects.

### Immunoblotting

Cells lysis was performed in 1.5×Laemmli buffer, and protein concentrations were quantified with the Pierce BCA Protein Assay from (Thermo Fisher Scientific Inc, MA, USA). Protein extracts (15 µg) were subjected to electrophoresis on either 10% or 12% sodium dodecyl-sulfate-polyacrylamide (SDS) gels and subsequently transferred to polyvinylidene difluoride membranes (PVDF, Merck Millipore, Burlington, MA, USA). The membranes were blocked with 5% milk in phosphate-buffered saline (PBS) and incubated with anti-human primary antibodies, followed by incubation with the corresponding horseradish peroxidase (HRP)-conjugated secondary antibodies. Immunoreactive bands were detected with horseradish peroxidase anti-mouse or anti-rabbit antibodies (Promega, Madison, WI, USA) using the enhanced chemiluminescence (ECL) system (Merck Millipore, Burlington, MA, USA). Immunoblot analysis was performed using the Li-COR Odyssey Imaging System (Lincoln, NE, USA) and Fusion Fx7 absolute (Vilber, Collégien, France).

### Flow cytometry

A total of 10,000 events per sample were analyzed using a BD FACSMelody cytometer (Becton Dickinson (BD) Biosciences, NJ, USA), and data was processed using the FlowJo software version vX.0.7 (Ashland, OR, USA). Cells were cultured in 6-well plates, in triplicate for each condition, and maintained at 37 °C in a 5% CO_2_ atmosphere with their respective media. Cell death was assessed using propidium iodide (PI; Invitrogen, Thermo Fisher Scientific Inc, MA, USA). Both floating and adherent cells were collected, centrifuged, and then resuspended in FACS buffer (PBS, 0.2% bovine serum albumin (BSA), and 2 mM EDTA), with 2 µg/mL of PI added just before analysis.

### Intracellular Fe^2+^ measurement

Ferrous iron was measured using FerroOrange (Dojindo Molecular Technologies, Kumamoto, Japan). After the experiment, cells were washed twice with PBS and resuspended in DMEM media without iron (Thermo Fisher Scientific Inc, MA, USA) supplemented with the FerroOrange probe (1 µM) for 30 min at 37 °C in a 5% CO_2_ environment, protected from light. Fluorescence intensity has been measured with a Xenius UVMC spectrofluorometer (SAFAS, Monaco). Fluorescence is normalized based on cell numbers of each condition.

### Immunofluoresence

DAOY WT and HD-MB03 WT cells were seeded on glass coverslips in 12-well plates at a density of 10,000 and 20,000 cells per well, respectively. The following day, cells were treated with 1 µM MMRi62 for 48 h or 100 µM deferoxamine (DFO) (Sigma D9533) for 24 h. At the end of the treatment period, cells were fixed with 4% paraformaldehyde (PFA) for 20 min at room temperature and washed three times with phosphate-buffered saline (PBS). Cells were subsequently permeabilized with PBS containing 0.1% Triton X-100 for 20 min and blocked with PBS containing 0.1% Triton X-100 and 1% bovine serum albumin (BSA) for 20 min at room temperature. Cells were then incubated overnight at 4°C with an anti-ferritin heavy chain (FTH) primary antibody (0.5 µg/mL; Invitrogen, 701934). The following day, coverslips were washed three times with PBS and incubated with the corresponding fluorescent secondary antibody (Cell Signaling Technology, 4412S anti-rabbit Alexa Fluor 488) diluted 1:500 in blocking buffer for 2 h at room temperature in the dark. Coverslips were subsequently washed three times with PBS and incubated with fluorescently labelled phalloidin (1:500; Cytoskeleton, PHDH1-A) for 1 h at room temperature to visualize the actin cytoskeleton. Following three additional washes with PBS, coverslips were mounted onto microscope slides using mounting medium containing DAPI (Vector Laboratories, Burlingame, CA, USA) for nuclear staining. Fluorescence images were acquired using a Zeiss fluorescence microscope (Carl Zeiss Microscopy GmbH, Jena, Germany) and processed using Zeiss imaging software (Carl Zeiss Microscopy GmbH, Jena, Germany).

### Generation, maintenance and transfection of cerebral organoids

Human induced pluripotent stem (iPS) cells (SCTi003-A, Stemcell Technologies, Vancouver, Canada) were maintained on a layer of Matrigel™ hESC-qualified (Corning, Thermo Fisher Scientific Inc, MA, USA), in mTeSR Plus medium (Stemcell Technologies, Vancouver, Canada). All cells were mycoplasma free. iPSC were dissociated with ReLeSR, an enzyme-free reagent without manual selection or scraping (Stemcell Technologies, Vancouver, Canada). Organoids were generated with the STEMdiff Cerebral Organoid Kit and the STEMdiff™ Cerebral Organoid Maturation kit (Stemcell Technologies, Vancouver, Canada) based on the protocol published by Lancaster et al. (17,18). At day 35 of differentiation, organoids were electroporated using a nucleofection protocol adapted from Denoth-Lippuner et al. (19). Briefly, five organoids were resuspended in 100 μL of Nucleofector Solution (Cell Line Nucleofector Kit V, Lonza) containing a total of 10 μg of plasmid : 2 μg of hyPBase (PiggyBac hyperactive transposase; pPB[Exp]-EGFP-CMV>hyPBase, VectorBuilder) together with 8 μg of PBCAG-mVenus for the control condition, or 2 μg of hyPBase together with 8 μg of PBCAG-MXmV, expressing c-Myc, OTX2, and mVenus. Electroporation was performed using program A-023 on a Nucleofector 2b device (Amaxa). Tumor development (presence and size) was monitored every 4 days using imaging (Macrofluo, Zeiss, Germany).

### UV–visible spectroscopy of MMRi62–Fe^3+^ interactions

The iron-chelating properties of MMRi62 were assessed by UV–visible absorption spectroscopy. In a quartz cuvette, a 28.3 μM MMRi62 solution in 4:1 MeOH / aqueous HEPES buffer (10 mM, pH 7.37) was treated with increasing amounts of a solution containing 1134 μM FeCl_3_·6H_2_O and 28.3 μM MMRi62 in MeOH. After each addition, the cuvette was shaken and absorption spectra immediately recorded over the 205–800 nm range at room temperature. To further evaluate the interaction of MMRi62 with Fe^3+^ under controlled iron availability, the experiment was repeated in the presence of nitrilotriacetic acid (NTA), a chelating agent that maintains Fe^3+^ in a soluble, ligand-bound form (20). In a quartz cuvette, a 28.3 μM MMRi62 solution in 4:1 MeOH / aqueous HEPES buffer (10 mM, pH 7.37) was treated with increasing amounts of a solution containing 609 μM FeCl_3_·6H_2_O, 670 μM NTA and 28.3 μM MMRi62 in MeOH. After each addition, the cuvette was shaken and absorption spectra immediately recorded over the 205–800 nm range at room temperature. Spectral changes upon Fe^3+^ addition were used to assess the interaction of MMRi62 with ferric iron. Analysis of the absorption spectra upon multicomponent analysis was developed using the Specfit software (21,22).

### Statistical analysis

In all figures, data are presented as the mean ± standard error of measurement (SEM) unless otherwise stated. Comparisons between multiple groups were performed using one-way or two-way analysis of variance (ANOVA) followed by Tukeys post hoc test as appropriate.

## Results

### MMRi62 triggers apoptotic cell death in medulloblastoma cells, coinciding with marked ferritin loss through autophagic and proteasomal pathways

Treatment of DAOY and HD-MB03 medulloblastoma cells with 1 or 10 μM MMRi62 resulted in a marked reduction in ferritin heavy (FTH) and light (FTL) chain protein levels. A concomitant decrease in NCOA4 expression was also observed in cells treated with 10 μM, suggesting an alteration of ferritin turnover pathways. Consistent with its reported activity on MDM2–MDM4 signaling (14,15) MMRi62 induced phosphorylation of p53 without significantly affecting total p53 protein expression (Figure 1A). To determine whether ferritin loss resulted from enhanced protein degradation, cells were co-treated with either the autophagy inhibitor bafilomycin A1 (Baf-1) or the proteasome inhibitor MG132. In both cell lines, inhibition of either pathway restored FTH and FTL levels, indicating that ferritin degradation was observed through both autophagic and proteasomal mechanisms upon MMRi62 treatment. Interestingly, p53 activation was maintained even in presence of Baf-1 and MG132 (Figure 1B). Functionally, 1 and 10 μM MMRi62 induced similar and strong reduction in cell viability after 48 hours of treatment in both DAOY and HD-MB03 cells. This cytotoxic effect was fully rescued by the pan-caspase inhibitor Q-VD-OPh, whereas the ferroptosis inhibitor ferrostatin-1 failed to provide protection, demonstrating that MMRi62 primarily induces apoptotic cell death under these conditions (Figure 1C). Furthermore, Baf-1 and MG132 had no effect on MMRi62-induced cell death (data not shown). Together, these findings demonstrate that MMRi62 induces apoptotic cell death in medulloblastoma cells, correlated with both autophagic and proteasomal ferritin degradation. This raises the possibility that ferritin turnover contributes to MMRi62 cytotoxicity.

**Figure 1.**
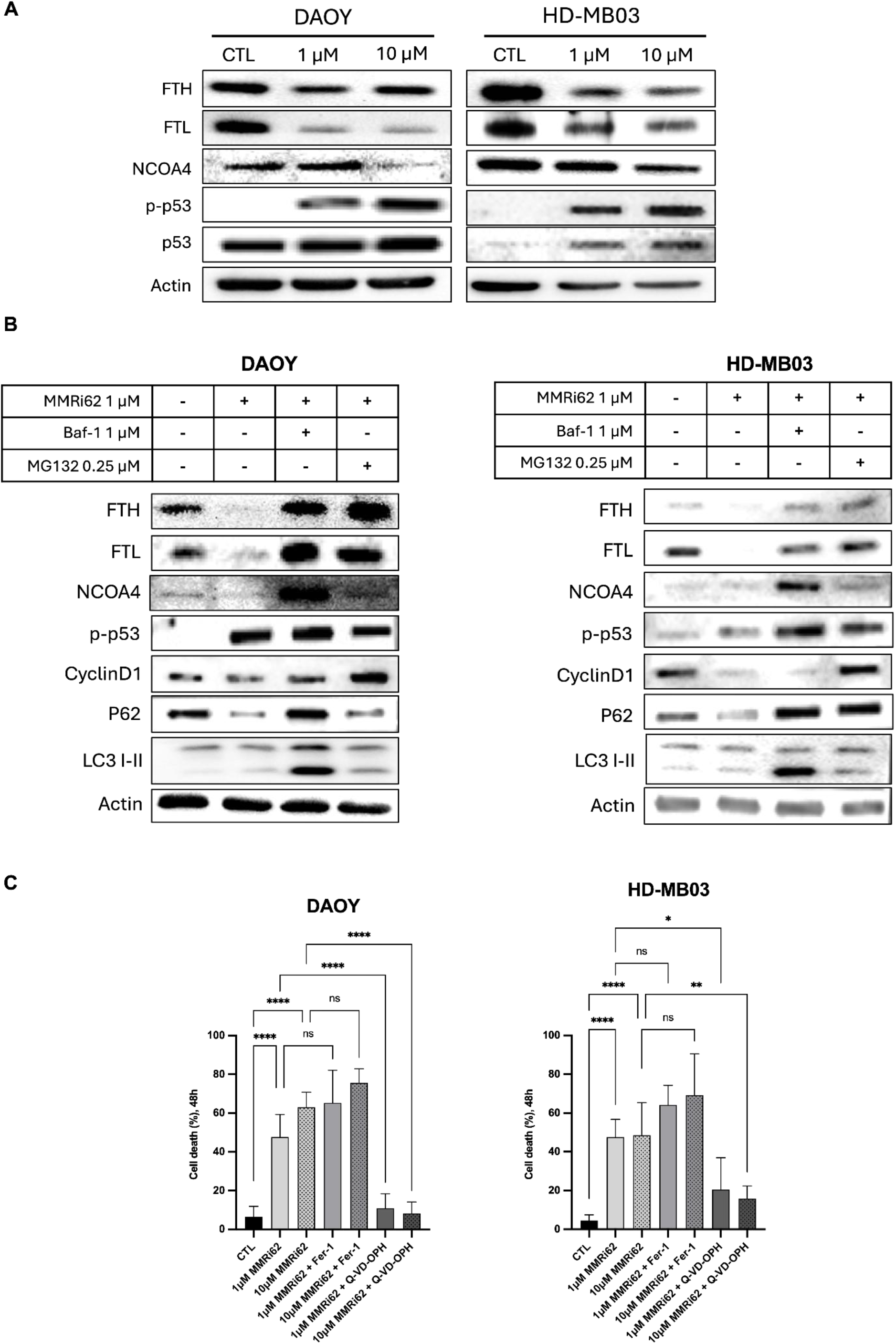
MMRi62 induces ferritin loss through autophagic and proteasomal pathways and triggers apoptotic cell death in MB cells. (A) Western blot analysis of DAOY and HD-MB03 cells treated with 1 or 10 µM MMRi62 for 24h. (B) DAOY and HD-MB03 cells were treated with 1 µM MMRi62 for 24h in the presence or absence of 1 µM Baf-1 or 0.25 µM MG132. Protein expressions of the markers of iron metabolism, autophagy or proteasomal activity were assessed by immunoblotting. (C) Cell viability of DAOY and HD-MB03 cells treated with 1 or 10 µM MMRi62 for 48 h was assessed using PI probe in the presence or absence of Fer-1, a ferroptosis inhibitor, or Q-VD-OPh, a pan-caspase inhibitor used to inhibit apoptosis. *Bar graphs represent the mean ± SEM (n = 3). Western blots and histograms are representative of three independent experiments. For WB analysis actin was used as a loading control*. ^*\**^*P<0.05*, ^*\*\**^*P<0.01*, ^*\*\*\**^*P<0.001*, ^*\*\*\*\**^*P<0.001. Abbreviations used: Medulloblastoma (MB), propidium iodide (PI), ferritin heavy chain (FTH), ferritin light chain (FTL), nuclear receptor coactivator 4 (NCOA4), tumor protein p53 (p53), Bafilomycin-A1 (Baf-1), sequestosoma-1 (p62), microtubule-associated protein 1 light chain 3 (LC3), ferrostatin-1 (Fer-1)*.

### -induced cell death occurs independently of ferritin degradation

Given that MMRi62 induces apoptotic cell death associated with ferritin loss, we next sought to determine whether ferritin degradation directly contributes to its cytotoxic activity. To address this question, we assessed the response to MMRi62 in ferritin-deficient models. Previously obtained, DAOY and HD-MB03 cells harboring knockout of *FTH1* (FTH KO cells) (9) displayed a similar reduction in viability compared to WT cells following treatment. In both cases, cell death was effectively rescued by Q-VD-OPh, indicating that apoptosis induction occurs independently of ferritin degradation (Figure 2A). As cell death was equivalent between WT and FTH KO cells, both in terms of magnitude and timing, we next investigated whether there were any differences in the cellular response to treatment. Kinetic analyses at 3, 6, 9, and 24 hours revealed that MMRi62 induced similar activation of p53 in both WT and FTH KO cells. In parallel, MMRi62 triggered a remodeling of iron metabolism markers, characterized by decreased ferritin levels, reflected by reduced FTL expression in FTH KO cells and reduced FTH and FTL in WT, and gradual increased expression of iron regulatory protein 2 (IRP2) and transferrin receptor 1 (TFRC1) (Figure 2B). Interestingly, this trend characteristic of iron-starvation response, was consistently observed across different MB cell lines, irrespectively of the culture medium used, as well as in the U87 glioblastoma cell line, in a time-dependent manner (Supp Figure 1), suggesting that this response reflects a broader and conserved cellular phenomenon rather than a cell line- or culture condition-specific effect. To further assess whether ferritin degradation contributes to MMRi62-induced cell death, we used NCOA4-deficient DAOY cells, in which ferritinophagy is impaired. Treatment with 1 μM MMRi62 induced a comparable cell death response in NCOA4-deficient and WT cells, indicating that MMRi62-induced cytotoxicity is independent of ferritin presence or its NCOA4-mediated degradation (Figure 2C). To further investigate this hypothesis, we analyzed ferritin protein levels in WT and NCOA4 KO cells treated with 1 μM MMRi62 for 24 h. Cells were also cultured in the presence or absence of Baf-1 and MG132 for WT cells, whereas NCOA4 KO cells were treated only with MG132, as these cells already exhibit impaired ferritinophagy due to disruption of the NCOA4-dependent autophagic pathway (11,23). As expected, NCOA4 KO cells show higher basal level of FTH and FTL protein. Interestingly, ferritin expression increased in NCOA4 KO cells following MMRi62 treatment. This accumulation, which was not observed in WT cells, suggests that NCOA4-mediated ferritin degradation is highly active in these cells upon MMRi62 exposure. As observed with WT and FTH KO cells previously (Figure 2B), sustained activation of p53 was observed upon treatment (Figure 2D). Together, these findings demonstrate that neither ferritin expression nor degradation is required for MMRi62-induced apoptosis and suggest that MMRi62 effect may instead alter iron homeostasis through a broader mechanism independent of ferritin degradation.

**Figure 2.**
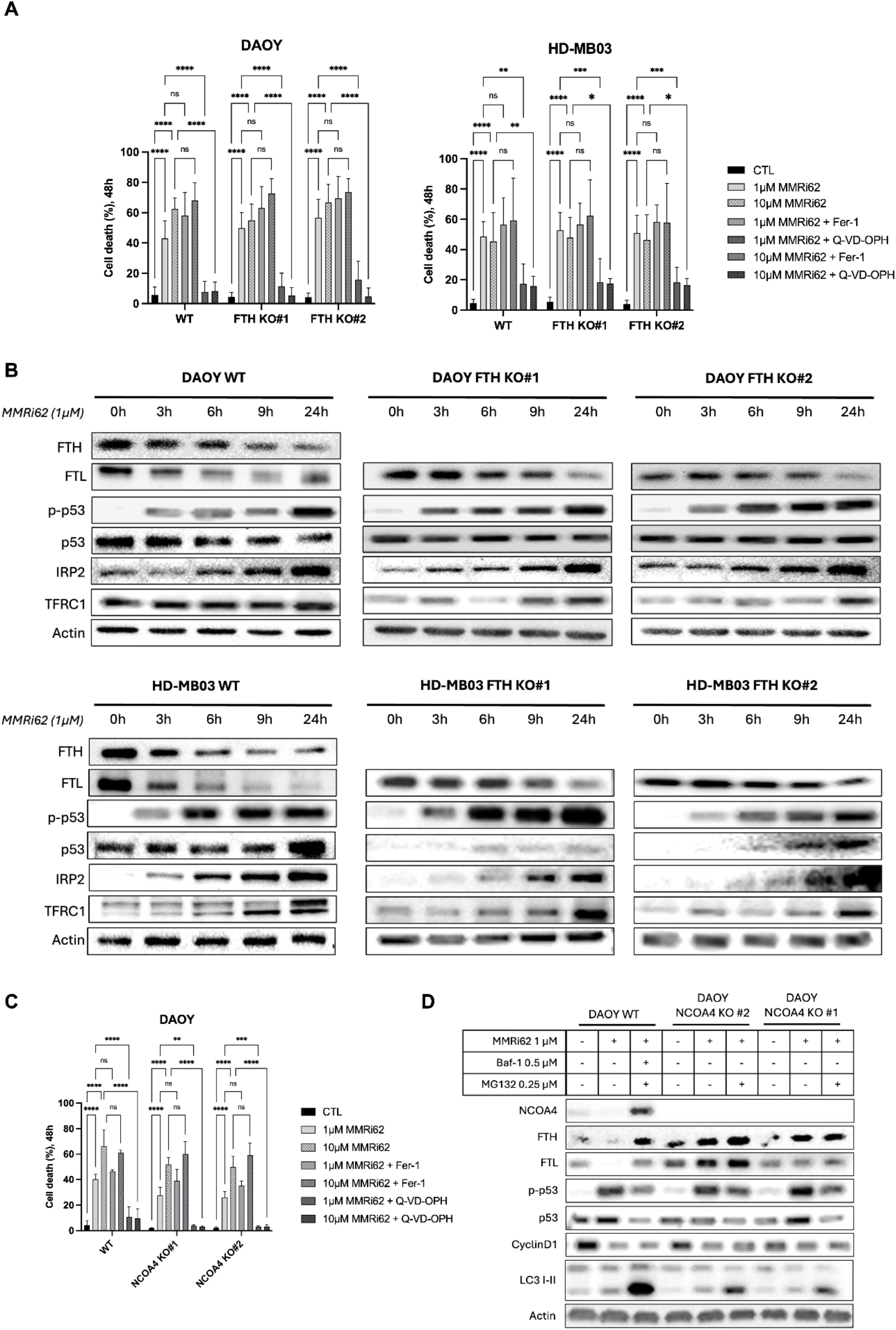
MMRi62-induced cell death is independent of ferritin degradation. (A) Cell viability of DAOY WT and FTH KO clones (#1 and #2), as well as HD-MB03 WT and FTH KO clones (#1 and #2), was assessed after 48 h of treatment with 1 or 10 µM MMRi62 in the presence or absence of Fer-1 or Q-VD-OPh. (B) A kinetic analysis of the expression of iron metabolism markers was performed by Western blotting after 3, 6, 9, and 24 h of MMRi62 treatment in DAOY and HD-MB03 WT and FTH KO cells. (C) The sensitivity of DAOY NCOA4 KO cells to 1 or 10 µM MMRi62 was assessed after 48 h of treatment in the presence or absence of Fer-1 or Q-VD-OPh. (D) Western blot analysis of DAOY WT and NCOA4 KO cells treated with 1 µM MMRi62 for 24 h in the presence or absence of 0.5 µM Baf-1 (WT cells only) or 0.25 µM MG132. *Bar graphs represent the mean ± SEM (n = 3). Western blots and histograms are representative of three independent experiments. For WB analysis actin was used as a loading control*. ^*\**^*P<0.05*, ^*\*\**^*P<0.01*, ^*\*\*\**^*P<0.001*, ^*\*\*\*\**^*P<0.001. Abbreviations used: ferritin heavy chain (FTH), ferritin light chain (FTL), nuclear receptor coactivator 4 (NCOA4), tumor protein p53 (p53), Bafilomycin-A1 (Baf-1), sequestosoma-1 (p62), microtubule-associated protein 1 light chain 3 (LC3), iron regulatory protein 2 (IRP2), transferrin receptor 1 (TFRC1), ferrostatin-1 (Fer-1)*.

### MMRi62 induces a cellular iron starvation response consistent with chelation-like mechanism

Given the coordinated changes observed in key iron homeostasis regulators, including increased IRP2 and TFRC1 expression together with reduced ferritin levels in WT and FTH KO cells (Figure 2B), we hypothesized that MMRi62 treatment decreases intracellular iron availability. Such iron depletion could activate adaptive iron-starvation pathways, thereby explaining the observed remodeling of iron metabolism in both DAOY and HD-MB03 cells. To test this hypothesis, we quantified intracellular ferrous iron (Fe^2+^) levels using FerroOrange following MMRi62 treatment. Analysis revealed a significant decrease in intracellular Fe^2+^ levels after 24 h of treatment with 1 μM MMRi62 in both DAOY and HD-MB03 WT cells (Figure 3A). Consistent with these findings, immunofluorescence analysis revealed a marked reduction in ferritin staining in MMRi62-treated cells, closely resembling the phenotype induced by the iron chelator deferoxamine (DFO) (Figure 3B,C). To further investigate the physiological relevance of this response, medulloblastoma-like tumours were genetically induced in brain organoids using a polycistronic PiggyBAC transposon coding for c-Myc, OTX2 and the fluorescent reporter gene mVenus as previously described (MXmV) (24). Tumors were made detectable by the expression of mVenus in neoplastic tissue, and organoids bearing tumors were treated with either MMRi62 or DFO. MMRi62 treatment progressively reduced the fluorescent tumor cell burden throughout the organoids, mirroring the effect observed with DFO treatment (Figure 3D). Western blot analysis of residual organoid revealed a common iron-starvation signature characterized by reduced ferritin expression and increased TFRC1 and IRP2 levels in MMRi62-treated organoids, closely resembling the molecular changes induced by the iron chelator, DFO (Figure 3E) and comparable to what we had observed in MB cell lines (Figure 2B). In addition, decreased levels of both OTX2 and c-MYC expressions were observed compared to their specific control in MXmV tumoral organoids treated with MMRi62 and DFO (Figure 3E). Given the 8-hydroxyquinoline-containing structure of MMRi62, a chemical motif previously associated with metal-chelating properties (25,26), together with the observation above, we hypothesized that MMRi62 may directly interfere with iron availability, acting as an iron chelator. This hypothesis was further supported by iron measurements performed under cell-free conditions. Incubation of MMRi62 in a cell-free system resulted in a reduction in detectable Fe^2+^ levels, again, comparable to the decrease in iron detection observed with DFO (Figure 3F).

**Figure 3.**
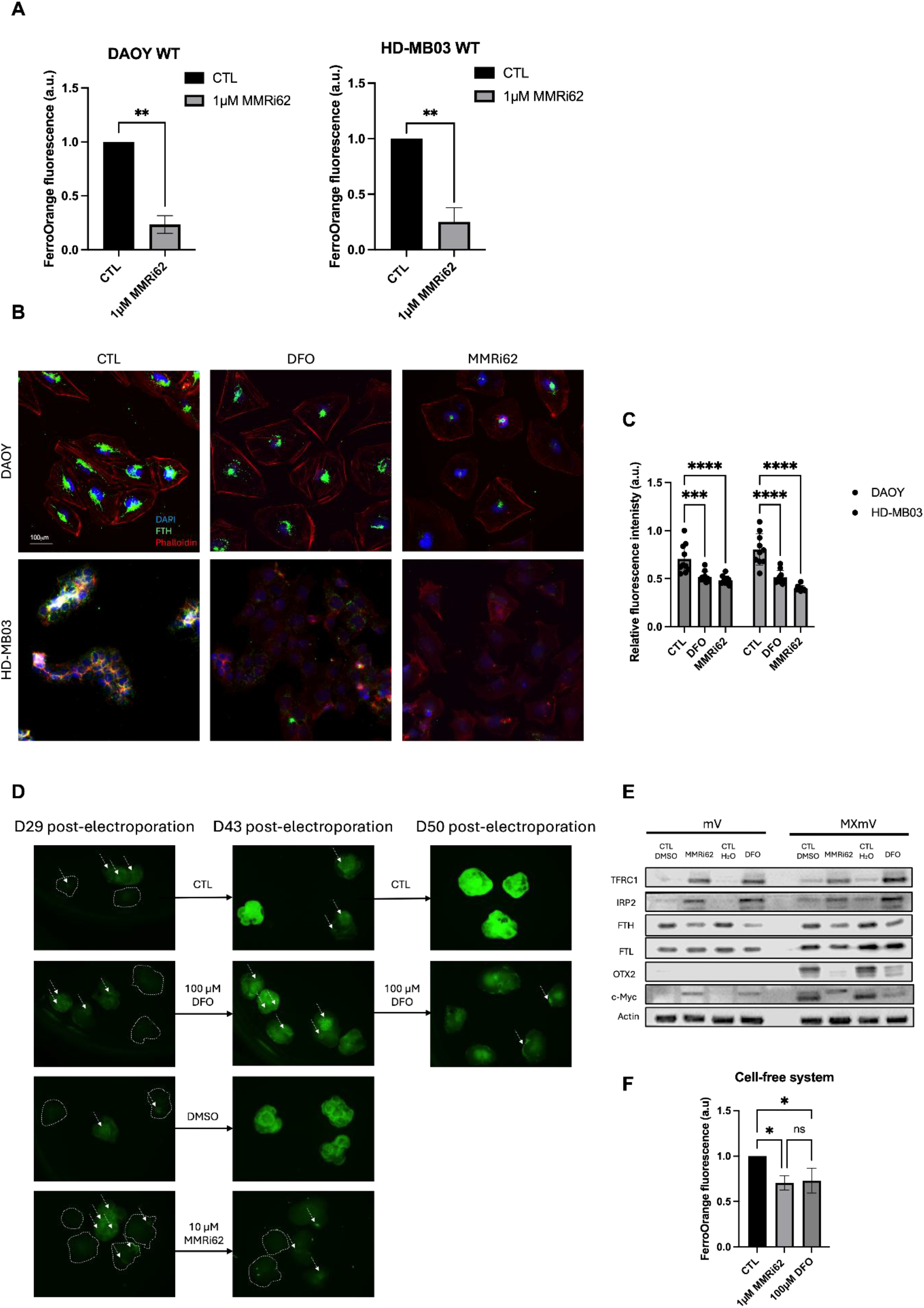
MMRi62 induces a cellular iron starvation response consistent with chelation-like mechanism. (A) Intracellular ferrous iron (Fe^2+^) levels in DAOY and HD-MB03 cells after 24 h of treatment with 1 µM MMRi62 using the FerroOrange probe. (B) Immunofluorescence staining of FTH (green) in DAOY and HD-MB03 WT cells treated or not with 1 µM MMRi62 for 48 h or 100 µM DFO for 24 h. Nuclei were stained with DAPI (blue) and the actin cytoskeleton with Alexa555-Phalloidin (red). (C) Quantification of FTH relative fluorescence intensity. (D) Evaluation of the effects of MMRi62 and DFO treatment on c-Myc/OTX2-driven medulloblastoma-like tumours genetically induced in brain organoids. Co-expression of the green, fluorescent mVenus reporter along with c-Myc and OTX2 (MXmV) allows the visualisation of tumor. Organoids were treated with 10 µM MMRi62 for 13 days or 100 µM DFO for 20 days. Tumor development was monitored every 4 days by fluorescence imaging. Organoids mV represents control organoid not transfected with OTX2/c-Myc. (E) Immunoblot analysis was performed at the end of the treatment period on control organoids expressing mVenus alone (mV) and tumoral MXmV organoids treated with vehicle control (CTL/DMSO and CTL/H_2_O), MMRi62 or DFO. The expression levels of TFRC1, IRP2, FTH, FTL, OTX2, and c-Myc were assessed. Actin was used as a loading control. (F) Free iron measurement (Fe^2+^) using FerroOrange in cell-free system with presence of 1 µM of MMRi62 or 100 µM of DFO. *Images, blots and histograms are representative of three independent experiments. Bar graphs represent the mean ± SEM (n = 3). Immunofluorescence images are shown at 40x magnification, with a scale bar representing 100* µm. *Western blots and histograms are representative of three independent experiments. For WB analysis actin was used as a loading control*. ^*\**^*P<0.05*, ^*\*\**^*P<0.01*, ^*\*\*\**^*P<0.001*, ^*\*\*\*\**^*P<0.001Abbreviations used: ferritin heavy chain (FTH), ferritin light chain (FTL), iron regulatory protein 2 (IRP2), transferrin receptor 1 (TFRC1), ferrostatin-1 (Fer-1), orthodenticle homeobox 2 (OTX2), cellular myelocytomatosis oncogene (c-Myc), deferoxamine (DFO), dimethyl sulfoxide (DMSO)*.

### MMRi62 interacts with Fe3+ and exhibit iron-chelating properties

To investigate whether MMRi62 interacts with iron, we monitored changes in its UV–visible absorption spectrum upon increasing concentrations of Fe^3+^. In our initial attempt, MMRi62 (28.3 μM) was titrated with increasing amounts of Fe^3+^ in a mixture of aqueous HEPES buffer (10 mM, pH 7.37) and MeOH. While a clear evolution of the absorption band around 250 nm was observed, diagnostic signals for the formation of molecular aggregates (*i.e*. stepwise increase of absorption all over the range 400-800 mn) rapidly appeared for amounts of Fe^3+^ as low as ∼0.3 equivalents, therefore precluding any reliable determination of association constant between iron and MMRi62 (Supp Figure 2). We therefore repeated the experiment in the presence of NTA (31.2 μM), which maintains Fe^3+^ in a soluble, ligand-bound form and provides more controlled iron availability (20). Under these conditions, increasing Fe^3+^ concentrations again produced progressive changes in the MMRi62 absorption spectrum while maintaining low levels of aggregation, as revealed by the lack of significant evolution of absorption in the range 400-800 nm (Figure 4). Analysis of the evolution of the absorption spectra upon Fe3+ binding indicates the formation of a 1:2 complex with MMRIi62 (log β = 10.5). Collectively these results supported a direct interaction between MMRi62 and Fe^3+^, consistent with iron chelating activity of MMRi62

**Figure 4.**
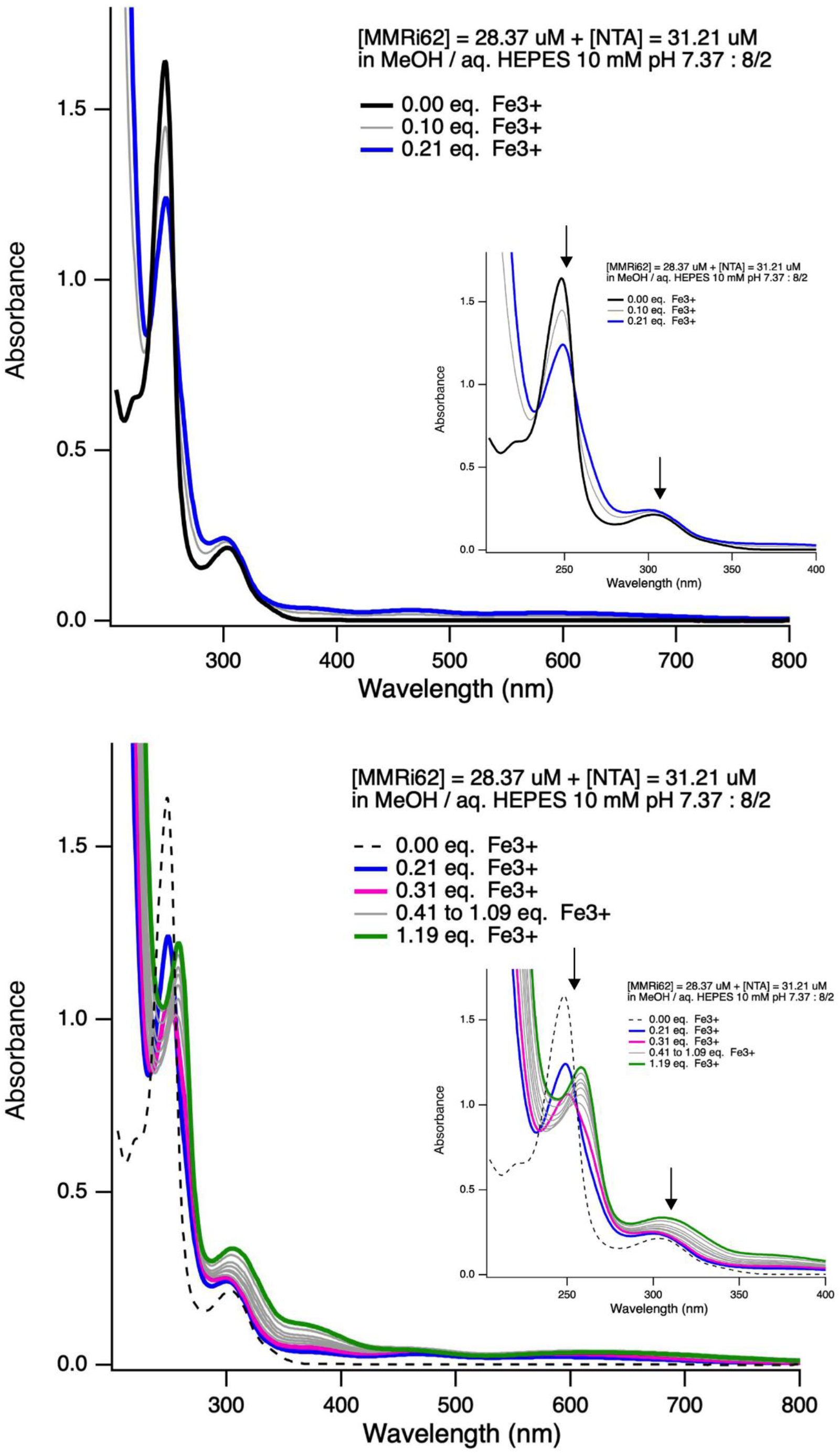
UV–visible spectroscopic analysis of MMRi62–Fe^3+^ interactions. UV–visible absorption spectra of MMRi62 (28.37 μM) in the presence of NTA (31.21 μM) and its evolution upon increasing concentrations of Fe^3+^ in MeOH/aqueous HEPES (10 mM, pH 7.37; 8:2). Upper panels show spectra obtained with 0, 0.10, and 0.21 equivalents of Fe^3+^ relative to MMRi62 over the 205–800 nm and 205–400 nm (insert) ranges. Lower panels show spectra obtained with increasing Fe^3+^ concentrations (0, 0.21, 0.31, 0.41–1.09, and 1.19 equivalents) over the same wavelength ranges. *Representative spectra are shown. Arrows indicate changes observed in the bands around 250nm pick and across the 260-320 nm region, consistent with Fe*^*3+*^ *dependent complex formation. Abbreviations used: MeOH (methanol), HEPES (4-(2-hydroxyethyl)-1-piperazineethanesulfonic acid), NTA (nitrilotriacetic acid)*.

## Discussion

Cancer cells exhibit profound alterations in iron metabolism to sustain their increased metabolic demands, while relying on ferritin to buffer iron and prevent iron-mediated toxicity. In medulloblastoma (MB), we previously identified ferritin as a potential therapeutic vulnerability (9), but the lack of pharmacological strategies to selectively target ferritin has limited its exploitation. Here, we investigated MMRi62 as a pharmacological approach to target ferritin and define the contribution of ferritin-dependent iron regulation to MB cell death.

Consistent with what have been reported previously (14), MMRi62 induced strong cytotoxic effect on MB cell lines. In addition, MMRi62 treatment reduced the population of c-MYC/OTX2-positive tumor-like cells (24), in a more physiologically relevant *ex vivo* cerebellar organoid model, demonstrating its cytotoxic activity against MB-like cells in a more complex tissue context and architecture. This cytotoxicity correlates with decreased expression of ferritin, mainly induced by enhanced degradation of ferritin with both proteasomal, which has already been proposed as a route for ferritin degradation (27) and canonical autophagic pathways in MB cell lines. However, contrary to the previously proposed ferroptotic mechanism, our findings demonstrate that MMRi62 induces predominantly apoptotic cell death. Indeed, MMRi62-induced cytotoxicity was efficiently rescued by the pan-caspase inhibitor (Q-VD-OPh), whereas pharmacological inhibition of ferroptosis failed to provide significant protection. These findings indicate that ferroptosis does not represent the primary mode of cell death induced by MMRi62 in our models and are consistent with previous reports describing apoptotic response observed with different MMRi compounds in other cancer contexts (15,28,29). MMRi compounds were initially identified as modulators of the MDM2/MDM4–p53 axis, with reported effects on MDM2 and/or MDM4 stability and subsequent activation of apoptotic signaling (13). Importantly, MMRi-induced cytotoxicity has also been reported in both p53-wild-type and p53-mutant settings, suggesting that their activity is not strictly dependent on functional p53 activity (28,29). Consistent with these observations, MMRi62 induced apoptosis in both HD-MB03 and DAOY cells despite their distinct p53 status. Thus, our findings place MMRi62-induced apoptosis within the broader cytotoxic activity of the MMRi family.

Beyond its effects on cell viability, MMRi62 markedly altered intracellular iron homeostasis. Treatment induced a significant reduction in intracellular Fe^2+^ levels, accompanied by remodeling of key iron-regulatory proteins, including IRP2 and TFRC1, in MB and glioblastoma cell lines, as well as MB-like organoid model. These findings indicate that MMRi62 alters intracellular iron homeostasis across distinct MB experimental models. These findings indicate that MMRi62 induces a broader remodeling of the cellular iron-handling machinery.

The observation that MMRi62 reduces ferritin expression in cancer cells initially appeared consistent with its previously proposed role as a ferritin destabilizer (14). However, our functional experiments demonstrate that ferritin degradation is not required for MMRi62- induced cytotoxicity. Using FTH deficient cells and NCOA4 deficient cells, in which NCOA4-dependent ferritinophagy is impaired, we demonstrate that neither ferritin expression nor its lysosomal degradation is necessary for MMRi62-induced apoptosis. These findings challenge the previously proposed model in which MMRi62 exerts its cytotoxic effects primarily through ferritin degradation and subsequent release of stored iron (14). Instead, the persistence of MMRi62-induced apoptosis in both *FTH1* and *NCOA4* deficient models indicate that ferritin downregulation occurs as a consequence of the primary effect of MMRi62. Observations from other members of the MMRi family further support the interpretation that ferritin degradation may occur as part of a broader MMRi-induced response rather than constituting a governing mechanism of cytotoxicity. For instance, MMRi71 has been reported to induce degradation of both MDM4 and FTH, while promoting robust apoptotic responses in leukemia models independently of p53 status (28). These observations demonstrate that ferritin degradation can occur in the context of MMRi-induced cytotoxicity without necessarily resulting in ferroptotic cell death. Although MMRi71 was not directly investigated in our study, the occurrence of ferritin degradation across different MMRi compounds suggests that modulation of ferritin stability may represent a recurrent feature of this compound class.

MMRi62 belongs to the quinolinol class of MMRi compounds and contains an 8-hydroxyquinoline scaffold, a chemical motif with well-established metal-chelating properties. 8-hydroxyquinoline and related derivatives have been extensively characterized for their ability to coordinate metal ions, including both Fe^2+^ and Fe^3+^, through the quinoline nitrogen and phenolic oxygen atoms (25,26). The presence of this metal-binding scaffold in MMRi62 therefore raises the possibility that its effects on intracellular iron homeostasis involve direct coordination and sequestration of iron. Consistent with this hypothesis, we demonstrated in the present study that MMRi62 directly interacts with Fe^3+^, providing experimental evidence that the compound can directly interact with iron ions and has the potential to sequester intracellular iron. In line with these observations, MMRi62 induced both cytotoxicity and alterations in iron homeostasis in cells, that closely resembled those observed with the classical iron chelator DFO. Both compounds reduced intracellular iron levels and affected cellular responses consistent with a state of iron deprivation, suggesting that MMRi62 may induce cellular effects, at least in part, through an iron-starvation-like mechanism. Importantly, our spectrophotometric analyses provide further support for a direct interaction between MMRi62 and iron. Addition of Fe^3+^ to MMRi62 produced concentration-dependent changes in its UV–visible absorption spectrum, and these changes were maintained in the presence of NTA, which limits the contribution of free Fe^3+^ by maintaining iron in a soluble, chelated form. Together with the cellular iron depletion and iron-starvation response observed following MMRi62 treatment, these findings support a model in which the 8-hydroxyquinoline (25,26) moiety of MMRi62 contributes to iron sequestration and thereby alters iron availability within MB cells.

Collectively, our findings redefine the biological activity of MMRi62 in medulloblastoma. Rather than acting primarily as a ferritin-targeting or ferroptosis-inducing compound, MMRi62 induces apoptotic cell death while profoundly perturbing intracellular iron homeostasis. The reduction in intracellular Fe^2+^, together with the remodeling of iron-regulatory pathways observed in both medulloblastoma cells and cerebellar organoids, indicates that alteration of iron availability is a prominent feature of MMRi62 activity. The 8-hydroxyquinoline scaffold of MMRi62, its established metal-chelating properties, and its ability to interact with Fe^3+^ provide a plausible mechanistic basis for this phenotype and raise the possibility that small molecule MMRi62 may act as an intracellular iron sequestrant. Importantly, these findings distinguish pharmacological modulation of iron homeostasis by MMRi62 from direct targeting of ferritin itself.

## Acknowledgments

The authors are sincerely grateful to all the associations, donors, and individuals whose generous financial contributions and continued encouragement made this work possible. We particularly thank the Fondation Flavien, Un Nouvel Espoir; the Fédération Enfants Cancers Santé; the Groupement des Entreprises Monégasques dans la Lutte contre le Cancer (GEMLUC); and the Savchuk Foundation for their longstanding and unwavering support.

## Author Approvals

All authors have read and approved the final version of the manuscript for submission to bioRxiv. This manuscript has not been published or accepted for publication elsewhere.

