## Supplementary Figures for "MMRi62 induces iron depletion–driven apoptosis through a ferritin-independent mechanism"

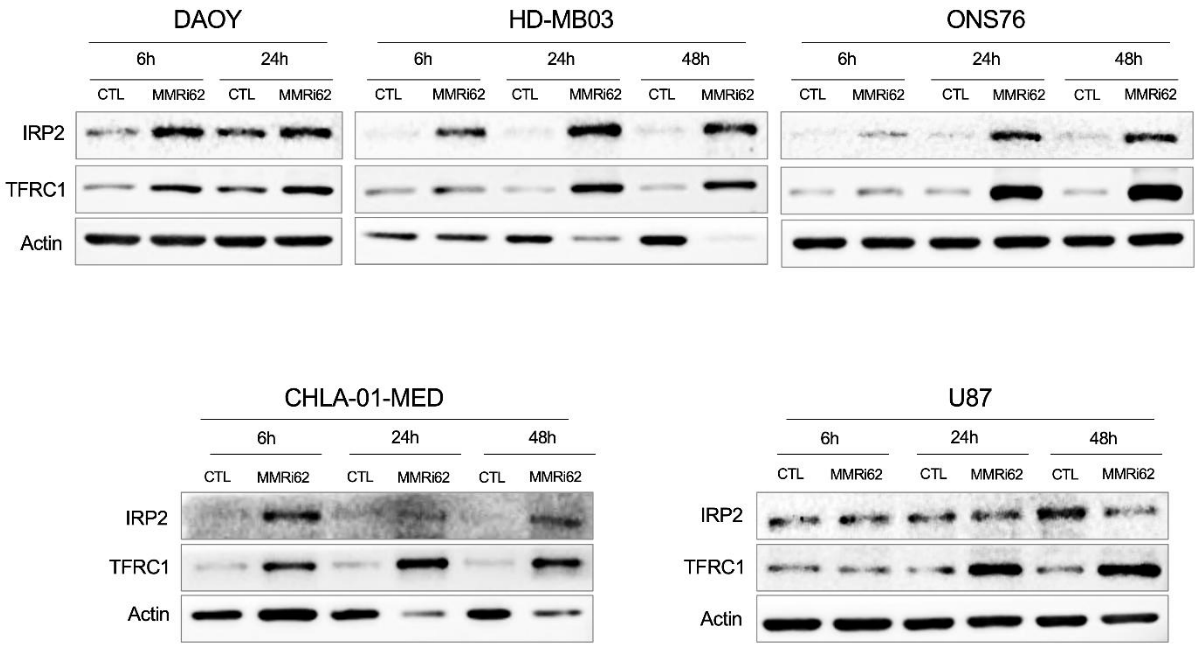


**Supplementary Figure 1 (related to Figure 2)**
A kinetic analysis of iron metabolism marker expression was performed by Western blotting following 24, 48, and 72 h of treatment with 10 µM MMRi62 in DAOY and HD-MB03 cells, as well as in two additional MB cell lines, ONS76 (cultured in the same medium as DAOY and HD-MB03) and CHLA-01-MED (cultured in defined stem-cell-enriching medium). Furthermore, the same kinetic analysis was performed in the representative glioblastoma cell line U87.

*Representative blots are shown.*
*Abbreviations used: iron regulatory protein 2 (IRP2), transferrin receptor 1 (TFRC1).*
*NB Cell death was observed in the case of all cell lines after 24h, and in the case of DAOY massive cell death at 48h time point prevented analysis.*

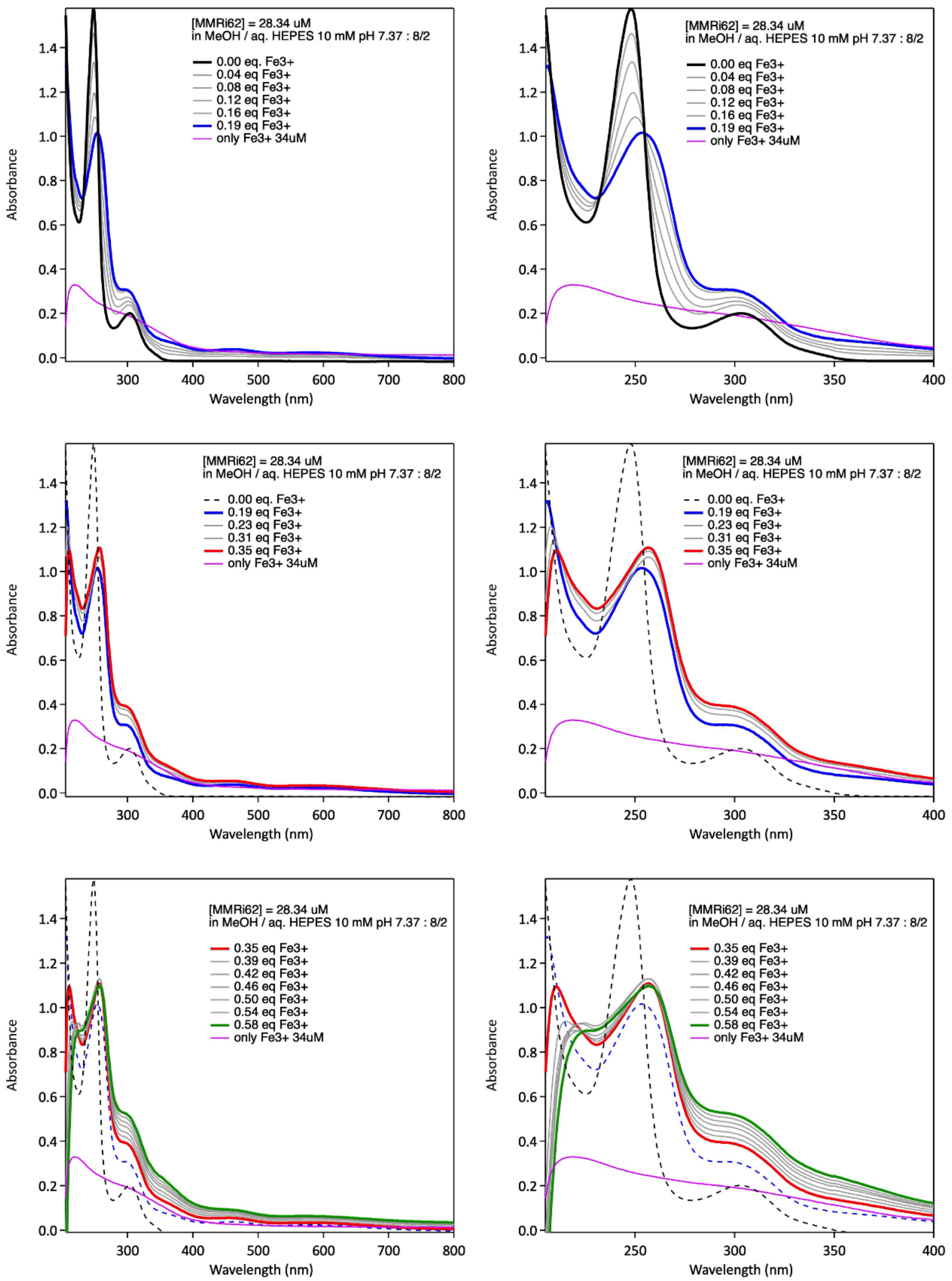


**Supplementary Figure 2 (related to Figure 4)**

UV–visible absorption spectra of MMRi62 (28.34 µM) and its evolution upon increasing concentrations of $Fe^{3+}$ in MeOH/aqueous HEPES (10 mM, pH 7.37; 8:2). Left panels show spectra recorded over the 205–800 nm range, while right panels show an expanded view of the 205–400 nm region. Spectra are shown for increasing $Fe^{3+}$ concentrations from 0 to 0.60 equivalents relative to MMRi62. The $Fe^{3+}$-only control (34 µM $Fe^{3+}$) is shown in magenta.

*Representative spectra are shown.*

*Abbreviations used: MeOH (methanol), HEPES (4-(2-hydroxyethyl)-1-piperazineethanesulfonic acid), NTA (nitrilotriacetic acid).*
